# RNA-seq analysis in seconds using GPUs

**DOI:** 10.64898/2026.03.04.709526

**Authors:** Páll Melsted, Elís Mar Guðnýjarson, Jóhannes Nordal

**Affiliations:** Faculty of Industrial Engineering, Mechanical Engineering and Computer Science, University of Iceland, Reykjavik, Iceland

## Abstract

We present a GPU implementation of kallisto for RNA-seq transcript quantification. By redesigning the core algorithms: pseudoalignment, equivalence class intersection, and the EM algorithm; for massively parallel execution on GPUs, we achieve a 30–50× speedup over multithreaded CPU kallisto. On a benchmark of 100 Geuvadis samples from Human cell lines the GPU version processes paired-end reads at a rate of 3.6 million per second, completing a typical sample in seconds rather than minutes. For a large dataset of 295 million reads, runtime drops from 40 minutes to 50 seconds. Our implementation demonstrates that careful algorithmic redesign, rather than naïve porting of software, is necessary to fully exploit the computing power of GPUs in sequence analysis.

## Introduction

The problem of transcript quantification from RNA-seq data has been studied extensively over the past decade. Initially, methods such as Cufflinks [20] and RSEM [13] focused on quantifying transcript abundance using RNA-seq reads aligned to a reference genome and an annotated transcriptome, with the alignment step relegated to specialized software packages such as TopHat [19], HISAT2 [11] or STAR [6]. Later, pseudoalignment approaches, pioneered by kallisto [4], abandoned the step of a full blown sequence alignment and focused rather on the task of finding the set of transcripts compatible with each sequencing read. This step is computationally faster and captures a sufficient statistic necessary for transcript quantification. Algorithmic innovations and improved software implementations reduced the computational burden from 20 CPU hours to 5-10 minutes and opened the possibility of processing RNA-seq datasets on a laptop or freely available notebooks in the cloud.

In the meantime, graphics processing units (GPU) have accelerated the world of scientific computing enabling both better runtimes and lower energy requirements for processing large datasets [5, 14]. However, the impact on sequencing based algorithms has been limited. GPU based implementations of sequence alignment [9, 10, 17, 1], i.e. edit distance and homology search, have received considerable attention, but do not translate directly to an end-to-end application for analysing a large set of sequencing reads, e.g. for RNA-seq or mapping genomic DNA. The success stories have been proprietary reimplementation of key parts of bwa and minimap2 for GPU acceleration by Parabricks, and early research on using GPUs for short read alignment [16, 12, 15] has not had a significant impact on large scale processing.

In this paper we considered the question of how to accelerate the kallisto software using GPUs. Framing this merely as a software problem is the wrong approach, to achieve this goal we must take apart all the algorithms designed with CPUs in mind and reassemble them under a different mental model of how the GPU works. The goal of this manuscript is not just to report on the success of this port, which enables processing RNA-seq data in seconds, but to show in detail how the algorithms have to be reconsidered with respect to a new programming and computer organization paradigm.

**Figure 1.**
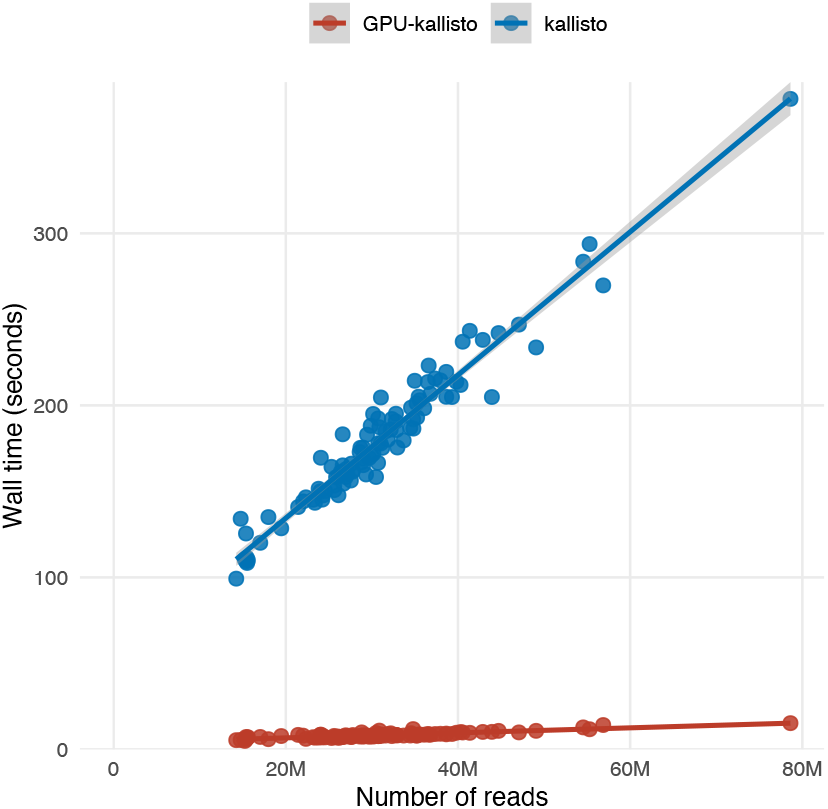
Benchmarking results for the GPU implementation of kallisto. The plot shows the time to process each of the 100 Geuvadis samples from Human RNA-seq, with GPU kallisto in red and CPU in blue as a function of the number of paired end reads in the sample.

## Results

To estimate the speed of the GPU implementation we ran kallisto and the GPU version on paired end reads from 100 samples of RNA-seq experiments from Human cell lines from the Geuvadis study [18]. Figure shows the benchmarking results, where the speedup for the GPU implementation is 30×, when setup time is excluded. We also ran both versions of kallisto on a large dataset of 295 million paired end reads where the runtime was 40 minutes for kallisto with 16 threads vs 50 seconds for the GPU implementation, a speedup of 48× . Table 1 shows an estimate of the components of the runtime when averaged across all 100 Geuvadis samples. We can see that the runtime is dominated by the EM algorithm and the I/O operations, even for bgzipped input, whereas the GPU mapping has a high throughput of processing 24.1M paired end reads per second.

**Table 1:** Breakdown of the runtime of components of GPU implementation of kallisto when averaged across all 100 Geuvadis samples.

| Component | CPU time | GPU time | GPU time per 1M paired end reads |
| --- | --- | --- | --- |
| Index Setup | 376 ms | 933 ms | - |
| I/O and decompression | 2370 ms | 841 ms | 27 ms |
| GPU mapping | 514 ms | 722 ms | 23 ms |
| EM algorithm | - | 3148 ms | - |

### The kallisto algorithm

Figure shows a schematic overview of the kallisto algorithm and how each step is implemented in the GPU framework. Each of the phases shown in the figure are described in the following sections.

Briefly the workflow of kallisto operates as follows. The transcriptome is given as a list of sequences referred to as targets or transcripts, *t*_1_, …, *t*_*T*_ . The algorithm operates on *k*-mers, sub-strings of a fixed length *k* from the transcripts. For each *k*-mer *x*, we define its equivalence class (EC) *e*_*x*_ as the set of all transcripts that contain the *k*-mer and we do not distinguish between a *k*-mer and its reverse complement.

To compute the equivalence class of a read we compute the intersection of all equivalence classes of the *k*-mers in the read,

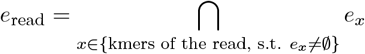

where *k*-mers in the reads that do not occur in the transcripts, i.e. the ones with an empty equivalence class, are not used in the intersection.

The equivalence class of each read is recorded and the counts of each equivalence class are a sufficient statistic to compute the abundance estimates of the targets.

The EM algorithm receives as input the count *c*_*e*_ of each equivalence class *e* as well as the effective length 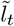 of each transcript *t*. We initialize a guess of the transcript abundance 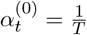 . In each step of the EM algorithm we alternate between computing 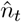 in the E step

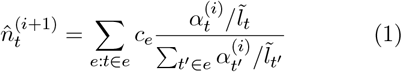

and normalizing in the M step

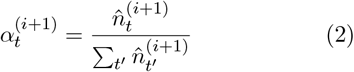

The algorithm iterates until convergence conditions are met.

kallisto stores an index for quick lookup of the equivalence class of a *k*-mer and stores new equivalence classes that occur as a result of the intersection that do not map to any one *k*-mer in the set of transcripts.

### EC lookup and intersection

Given a kallisto index we keep track of the following information on the GPU. For each equivalence class *e*_*i*_ we store the set of transcripts 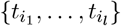 in sorted order as a flattened array *E* with offsets stored to the start of each equivalence class. The *k*-mer lookup is performed by a hash table, stored on the GPU implemented via cuCollections, mapping a *k*-mer *x* to the id of the equivalence class *e*_*x*_. The *k*-mers are stored as 64-bit integers via 2-bit encoding of nucleotide sequences. Additionally we keep track of the inverse map of the set of transcripts to their corresponding equivalence class ids in a dynamic hash table stored on the GPU.

For each read we process we know the offsets into the read buffer where the sequences reside. Using this information we generate all the *k*-mers *x* and perform a hash table lookup to compute *H* [*x*], all of which can be done in parallel. Computing the intersection of ECs for each read can be done in parallel as each read is independent. However, the size of the intersection is unknown, which complicates memory allocation. To solve this we break the task of taking a list of ECs for a read *e*_0_, …, *e*_*n*−1_and computing the intersection, as an ordered set of transcript ids as follows.

First the ECs are deduplicated, since adjacent *k*-mers are likely to map to the same equivalence class, and empty equivalence classes are removed. If more than one equivalence class remain we need to compute their intersection. The computation of the intersection finds the smallest EC and uses the size as an upper bound on the size of the output. We then iteratively compute the intersection, starting with the smallest EC, by either using a mergesort-like comparison when the sets are of a similar size or a binary search when one set is much smaller. For paired end reads we compute the intersection of each read separately before computing the final intersection.

The description above would suffice to implement this on a CPU, however we do not have the luxury of dynamic memory allocation per thread on the GPU for intermediate memory required when either deduplicating the ECs or computing the intersection of transcript lists. The solution to this is to have each thread use shared memory *x* where each thread *i* only uses segments *x* [*a*_*i*_ : *b*_*i*_]. To compute values for the upper and lower bounds we perform two-passes, in the first pass each thread estimates the amount of memory required *m*_*i*_. We then compute 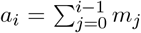 and *b*_*i*_ = *a*_*i*_ + *m*_*i*_, the *a*_*i*_ values are computed in parallel by a prefix-scan, a fundamental algorithm in parallel computing [3]. Once the memory boundaries have been determined each thread can operate on its segment without having to coordinate with other threads. In the case of the EC deduplication the memory required is the number of *k*-mers, whereas for the transcript set intersection we use the smallest EC as an upper bound on the intermediate memory required.

Finally when the intersection is available we look up the ordered set of transcript ids in a hashtable that maps the sets to their respective equivalence classes and update the equivalence class counts. When the intersection results in a set that is not found in the map we store the set and update the hashtable when we are done processing the batch of reads.

### The EM algorithm

Before iterating the EM steps we form a transpose index, for each transcript *t*, the set of equivalence classes *e* with *t* ∈*e*. To map the EM algorithm to the pipeline we break down equation 1 where the contribution of each EC count, *c*_*e*_ is divided up among the transcripts *t* ∈ *e* according to the expected probability of being the correct source, normalized by the effective length of the transcript 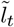 .

To compute this we first compute for each equivalence class, transcript pair (*e, t*) the term

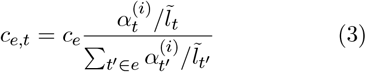

To do this on the GPU we first compute the de-ing the transposed index we compute the sum for nominator, 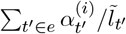, for each *e* in parallel. Us-each transcript *t* in parallel 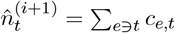 . With this strategy we can employ the entire GPU to compute a single E-step of the EM algorithm. The M-step is trivial, set 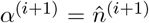 and for efficiency reasons we only check for convergence every 10 iterations.

**Figure 2.**
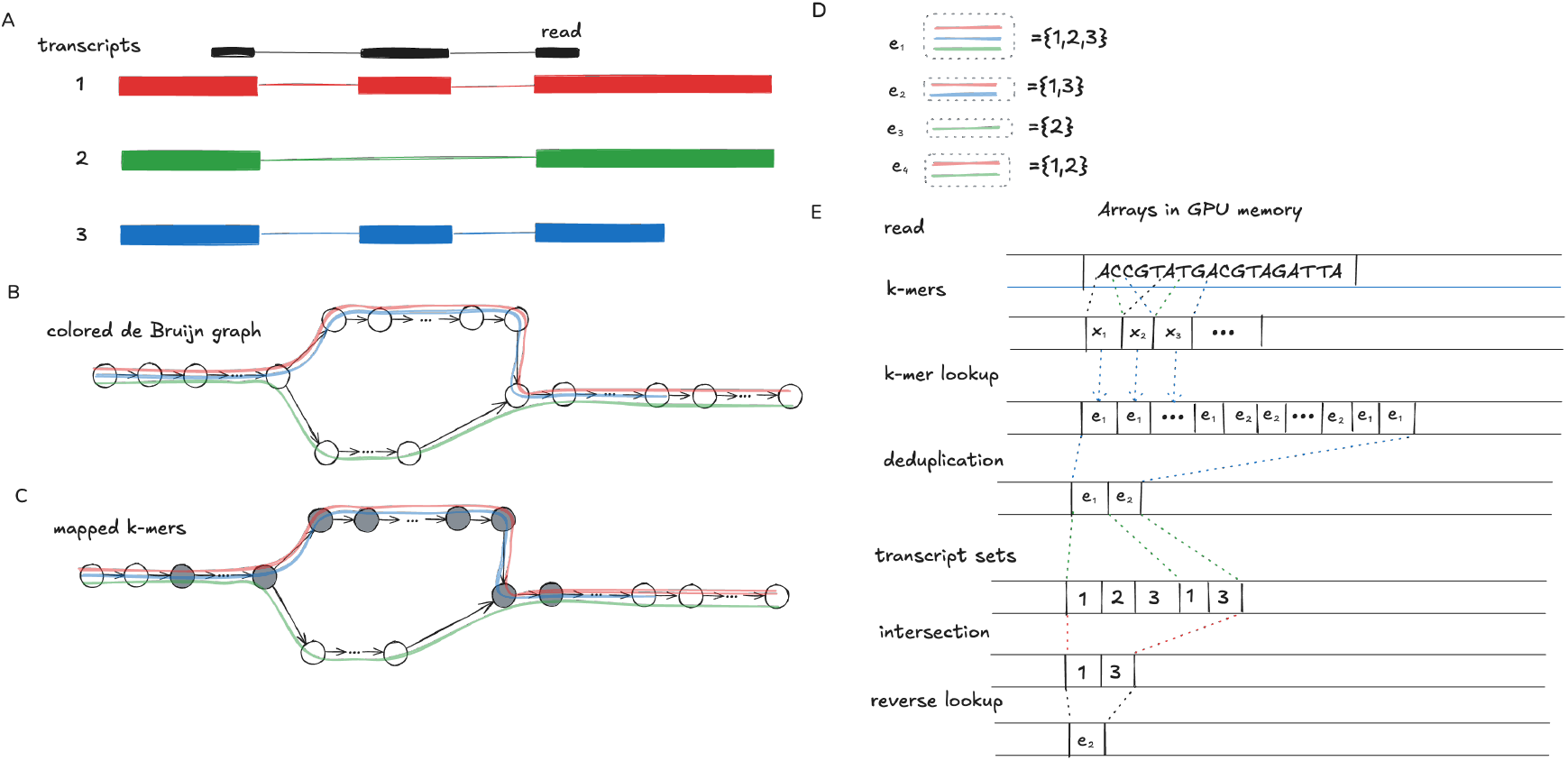
A schematic overview of the kallisto algorithm and how it each step is implemented in the GPU framework. **A** A schematic of three transcripts (colored) from one gene, sharing an exon structure and a sequencing read (black). **B** The target de Bruijn graph showing *k*-mers (nodes) and the paths traced by each transcript in the graph. **C** The *k*-mers in the sequencing read correspond to a path in the de Bruijn graph shown by shaded nodes. **D** The set of equivalence classes in the graph. **E** The processing of the read in GPU memory, read sequences are transformed to *k*-mers, each *k*-mer is mapped to an equivalence class, the set of equivalence classes are deduplicated, each equivalence class is expanded to its corresponding transcript set, the intersection of the transcript sets is computed, and the result is converted into the corresponding equivalence class.

### FASTQ parsing and decompression

Parsing a FASTQ file looks like a solved problem best left to established libraries such as the `kseq` package. However the enormous computing power of the GPU and the harsh reality of Amdahl’s law [2] leave parsing the input as a bottleneck dominating the runtime. Even worse, since most sequencing reads are compressed we need to decompress the file before parsing. While gzipped files can be decompressed in a streaming fashion they are inherently serial. To address this we split the decompression and parsing into two separate steps.

When the input is compressed with bgzip we load the file using the CPU, identify a suitable number of blocks and move to the GPU. Each block of the bgzip input is then decompressed on the GPU using the `nvcomp` library. For gzipped input we decompress the buffer on the CPU and move the resulting text buffer to the GPU.

Once the textbuffer resides in GPU memory we need to determine the boundaries of each read. To do this in parallel we launch kernels where each thread is given a segment to work with to identify the presence of newlines at any given position. Given the boolean array of newlines we compute the global location of each newline using a prefix-scan [3] operation. Since each record in the FASTQ file consists of 4 lines of text we identify where the reads are in the buffer in parallel.

## Discussion

We have presented a GPU implementation of the kallisto software for RNA-seq transcript quantification. The graphics processing unit used in the benchmark, GeForce RTX 5090, is a high-end consumer GPU and yet is powerful enough to accelerate the kallisto software by a factor 30-50× . Although we ran the benchmark on a GPU with the latest architecture, Blackwell, the code will run on NVIDIA GPU generations from Volta and later. Early on we discovered that the task of getting the sequences from a compressed FASTQ file to the GPU was a bottleneck. To solve this we relegated the task of decompression and parsing to the GPU, using parallel algorithms developed long before the advent of GPUs or the massively parallel CPUs of today.

This is a key challenge that the bioinformatics community must face in the future, parsing the data must not be a serial bottleneck, as it is with gzipped FASTQ files if we are to fully utilize the GPU resources. Just the simple act of producing bgzipped FASTQ files enables parallel processing and reduces the total time spent on the CPU decompressing the sequences. To emphasize this, for the largest files processed, just copying the files using `dd` took 25 seconds, whereas the running time of GPU kallisto was 50 seconds. When processing it by counting the lines `gzip -c | wc –l`took 10 minutes on the CPU.

The work presented here opens up several new avenues of research for the use of GPUs in bioinformatics and computational biology. Sequence analysis can be accelerated but thought must be put into getting the data on the GPU as soon as possible and algorithms have to be adapted to massively parallel processing where the convenience of dynamic memory allocation is not always an option. The data structures developed for read mapping, e.g. the FM-index [8], can perhaps be adapted to batch processing of reads in parallel, the limited memory on the GPU may make them an attractive option.

## Methods

### Datasets

One hundred samples from Geuvadis were downloaded from the SRA database and converted to bgzipped FASTQ files. Additionally we processed a dataset consisting of 295 million paired end reads (SRR30898520) from human RNA-seq data.

### Benchmarking

Benchmarks were run on a workstation with an AMD Ryzen 9 9900X (12 cores), 128 GB RAM, a 3.6 TB NVMe SSD (Samsung 990 PRO 4TB), and an NVIDIA GeForce RTX 5090 (32 GB RAM). Both kallisto and GPU-kallisto were compiled with the GCC compiler and the CUDA toolkit and run with default parameters, but provided the same mean effective length for both versions (`-l 175 -s 25`), determined by the mean of the effective lengths of the first 10 samples from the Geuvadis experiment. The index was constructed using the cDNA sequences from the Ensembl release 113 [7]. The kallisto benchmarks were run in parallel on all 12 cores of the CPU, each job utilizing 4 threads. The GPU benchmarks were run serially with 4 threads and a single GPU.

### Software and availability

The GPU version of kallisto uses the CUDA framework to run code on NVIDIA GPUs and uses the `cccl` library for prefix scan and several other parallel algorithms, the `nvcomp` library for decompression and `cuCollections` for the hash table implementations.

The code for the GPU implementation of kallisto is available on GitHub at https://github.com/pachterlab/kallisto under the `gpu` branch.

## Acknowledgements

The authors would like to thank Lior Pachter for helpful discussions and feedback on the manuscript. LLM models were used to assist with the optimizing and writing parts of the software, primarily Opus 4.5 and Opus 4.6 from Anthropic, all writing of the manuscript was performed by humans.

## Author contributions

Work on this paper was lead by P.M., who conceived of the project, implemented the software, analyzed the datasets and wrote the initial draft of the manuscript. P.M. and J.N. implemented the GPU code for mapping *k*-mers to equivalence classes and computation of the intersection of transcript sets. P.M. and E.M.G developed the GPU code for the EM algorithm. All authors contributed to the writing of the manuscript.

## Notes

### Competing Interest Statement

The authors have declared no competing interest.

